# Targeting SARS-CoV-2 infection through CAR-T like bispecific T cell engagers incorporating ACE2

**DOI:** 10.1101/2022.01.19.476940

**Authors:** Mikail Dogan, Lina Kozhaya, Lindsey Placek, Fatih Karabacak, Mesut Yigit, Derya Unutmaz

## Abstract

Despite advances in antibody treatments and vaccines, COVID-19 caused by SARS-CoV-2 infection remains a major health problem resulting in excessive morbidity and mortality and the emergence of new variants has reduced the effectiveness of current vaccines. Here, as a proof-of-concept we engineered primary CD8 T cells to express SARS-CoV-2 Spike protein-specific CARs, using extracellular region of ACE2, and demonstrated their highly specific and potent cytotoxicity towards Spike-expressing target cells. To improve on this concept as a potential therapeutic, we developed a bispecific T cell engager combining ACE2 with an anti-CD3 scFv (ACE2-Bite) to target infected cells and the virus. Similar to CAR-T cell approach, ACE2-Bite endowed cytotoxic cells to selectively kill Spike-expressing targets. Furthermore, ACE2-Bite neutralized the pseudoviruses of SARS-CoV, SARS-CoV-2 wild-type and variants including Delta and Omicron, as a decoy protein. Remarkably, ACE2-Bite molecule showed a higher binding and neutralization affinity to Delta and Omicron variants compared to SARS-CoV-2 wild-type Spike proteins, suggesting the potential of this approach as a variant-proof, therapeutic strategy for future SARS-CoV-2 variants, employing both humoral and cellular arms of the adaptive immune response.

## Introduction

Worldwide, Coronavirus disease 2019 (COVID-19) caused by SARS-CoV-2 infection has caused close to 6 million deaths and a chronic debilitating condition called Post-Acute COVID-19 Syndrome (PACS) in many millions more. An unprecedented effort by researchers around the world has resulted in the development of a spectrum of preventative and therapeutic approaches at an extraordinary speed. Vaccines focused on virus Spike protein (such as messenger RNA vaccines, non-replicating vector vaccines, and virus-like particle vaccines) are highly efficient in preventing infection (1). Several therapeutic developments, such as synthetic neutralizing antibodies, monoclonal antibodies to Spike protein (2) (3), (4, 5) and immunomodulators such as corticosteroids (6, 7), anti-IL-6 (8), anti-IL-1 (9) and Interferon-β-1a agents (10) were shown to have a range of treatment efficacy from non-effective to highly promising. Some of those treatments were repurposed to focus on blocking viral entry while others were used to control the hyperinflammatory immune response during the disease. Beyond antibody therapies, specific SARS-CoV-2 immunomodulators have not been developed yet are needed as specific antibodies potentially lose their effectiveness due to new variants. Therefore, the development of treatment approaches that remain effective against SARS-CoV-2 variants is of great interest.

SARS-CoV-2 uses its Spike protein to bind to the key host receptor Angiotensin-Converting Enzyme 2 (ACE2) on target cell surface for cell entry (11) and mutations in Spike protein result in higher affinity of the virus to ACE2 (12) and/or a better escaping mechanism from the immune system (13). Following the cell entry, SARS-CoV-2 generates viral components by taking over the protein synthesis machinery of the host cell and displays Spike protein on the cell membrane (14). Using ACE2 molecule to target these Spike-expressing infected cells could be an effective strategy in preventative and therapeutic approaches to COVID-19 in the future since ACE2 receptor would have to be compatible to the binding of forthcoming mutant Spike proteins.

Here, we used a synthetic biology approach to engineer primary human CD8 T cells to express Spike protein-specific chimeric antigen receptors with ACE2 or anti-Spike antibody (ACE2 CAR or anti-Spike CAR) on the extracellular domain to target SARS-CoV-2 infected cells. Considering a viable strategy in clinical setting, we also engineered a novel ACE2/anti-CD3 bispecific T cell engager antibody (ACE2-Bite) to target both SARS-CoV-2 infected cells and the SARS-CoV-2 virus. Bispecific T cell engager antibodies (Bites) are engineered chimeric molecules that are designed to bind to CD3 on T cells via an anti-CD3 single chain variable fragment (ScFv) and to a target cell via a target-specific molecule. Upon bridging the T cells with a target cell, Bites trigger T cell activation and subsequent target cell apoptosis. CAR-T and bispecific T cell engager antibody approaches we developed in our study are similar to methods targeting cancer cells, several of which have been approved for treatment (15).

In this study, we show that ACE2-CAR and anti-Spike CAR-expressing CD8 T cells became activated and selectively killed different types of target cells expressing SARS-CoV-2 Spike protein on their surface. The ACE2-Bite antibodies also led to T cell activation in the presence of Spike expressing targets, and mediated cytotoxicity to these targets. In addition, ACE2-Bite antibodies acted as a decoy receptor for pseudoviruses incorporated with Spike proteins of Coronaviridae including SARS-CoV, SARS-CoV-2 wild-type, Delta and Omicron variants and neutralized these with a significantly increased affinity to Delta and Omicron. Taken together, these results suggest that the novel chimeric antigen receptors and bispecific antibodies may be used to re-direct cytotoxic immune cells towards SARS-CoV-2 infected host cells and to neutralize variant strains of the virus.

## Materials and Methods

### ACE2 CAR construct

CAR constructs consisting of CD8 alpha signal peptide, extracellular domain of ACE2 molecule or single chain variable fragment (scFv) of anti-CD19 or anti-Spike protein antibodies, CD8 hinge domain, CD8 transmembrane domain, 4-1BB (CD137) intracellular domain and CD3ζ domain were designed with Snapgene and synthesized via Genscript. ACE2 extracellular domain, CD8a signal peptide, CD8 hinge, CD8 transmembrane domain, 4-1BB intracellular domain and CD3ζ domain sequences were obtained from Ensembl Gene Browser and codon optimized with SnapGene by removing the restriction enzyme recognition sites that are necessary for subsequent molecular cloning steps, while preserving the amino acid sequences. Anti-CD19 and anti-Spike scFv amino acid sequences were obtained from Addgene plasmids #79125 and #155364, respectively, reverse translated to DNA sequences and codon optimized with Snapgene 5.2.4. The constructs were then cloned into a lentiviral expression vector with a multiple cloning site separated from RFP reporter via an Internal Ribosomal Entry Site (IRES).

### Spike protein constructs

Human codon optimized wild-type and Omicron full-length SARS-CoV-2 Spike protein sequences were synthesized by MolecularCloud (MC_0101081 and MC_0101272, respectively) and then cloned into pLP/VSVG plasmid from Thermo Fisher under CMV promoter after removing the VSVG sequence via EcoRI-EcoRI restriction digestion. 5’-ACGACGGAATTCATGTTCGTCTTCCTGGTCCTG-3’ and 5’-ACGACGGAATTCTTAACAGCAGGAGCCACAGC-3’; and 5’-ACGACGGAATTCATGTTCGTGTTCCTGGTGCT-3’ and 5’-ACGACGGAATTCTTAACAGCAACTGCCGCAG-3’ primers were used to generate wild-type and Omicron SARS-CoV-2 Spike protein sequences without the Endoplasmic Reticulum Retention Signal (ERRS, last 19 amino acids of Spike) (14), respectively. For stable wild-type Spike protein overexpression, wild-type and Omicron Spike protein sequences without ERRS domain was cloned into a lentivector with a GFP marker under LTR promoter. Human codon optimized Delta full-length SARS-CoV-2 Spike protein plasmid was synthesized by Invivogen (plv-spike-v8).

### VSVG and Spike Protein pseudotyped lentivirus production

The lentiviruses pseudotyped with vesicular stomatitis virus G protein envelope were generated with HEK293T cells. Briefly, the lentivector plasmids containing the constructs were co-transfected with vesicular stomatitis virus G protein, pLP1, and pLP2 plasmids into HEK293T cells at 80–90% confluency using Lipofectamine 3000 (Invitrogen) according to the manufacturer’s protocol. In the case of Spike protein pseudotyped lentiviruses, a lentivector plasmid containing GFP reporter was co-transfected with wild-type or mutated SARS-CoV-2 Spike protein plasmids in the same manner. The transfection medium was replaced with RPMI 1640 with 10% FBS 6 hours post-transfection. Viral supernatants were collected 24 to 48 hours post-transfection and filtered through a 0.45-μm syringe filter (Millipore) to remove cellular debris. A Lenti-X concentrator (Takara Bio USA) was used according to the manufacturer’s protocol to concentrate the virus 10-20× and the resulting lentiviral stocks were aliquoted and stored at −80°C. To measure viral titers of VSV-G pseudotyped lentiviruses, virus preparations were serially diluted on Jurkat cells and 3 days post-infection, infected cells were measured using flow cytometry and the number of cells transduced with 1 mL of virus supernatant was calculated as infectious units per milliliter. For spike protein pseudotyped lentiviruses, to measure viral titers, virus preparations were serially diluted on ACE2 over-expressing 293 cells, which were stained for their ACE2 expressions and confirmed ∼%100 positive. 72 hours after infection, GFP positive cells were counted using flow cytometry and the number of cells transduced with virus supernatant was calculated as infectious units/per mL. Based on these titer values, primary T cells, 293 T cells and T2 cells were transduced with a multiplicity of infection (MOI) of 3–10.

### ACE2-Bite design and production

The ACE2-Bite construct consisting of ACE2 signal peptide, ACE2 extracellular domain, a linker peptide, an anti-CD3 antibody single-chain variable fragment, a His-Tag, and a Hemagglutinin (HA) Tag was designed with Snapgene and synthesized via Genscript. ACE2 signal peptide and extracellular domain sequences were obtained from Ensembl Gene Browser (Transcript ID: ENST00000252519.8). Anti-CD3 antibody single-chain variable fragment, His-Tag, and Hemagglutinin (HA) Tag sequences were obtained from Addgene plasmid #85437. ACE2-Bite construct was cloned into an RFP marked lentivector under LTR promoter, and Expi293F™ suspension 293 cells from ThermoFisher were transduced with the ACE2-Bite expressing VSVG pseudotyped lentiviruses with multiplicity of infection of 5. The cells were then grown in Expi293™ Expression Medium in shaking flasks for 7 days until they reached maximum viable density. ACE2-Bite containing supernatant was then collected and filtered/concentrated up to 30-fold with 30kDa MilliporeSigma™ Amicon™ Ultra Centrifugal Filter Units. Concentrated ACE2-Bite and control supernatants were aliquoted and stored in 4°C.

### Engineering CAR-T cells and Spike expressing target cells

Healthy adult blood was obtained from AllCells. PBMCs were isolated using Ficoll-paque plus (GE Health care). CD8 T cells were purified using Dynal CD8 Positive Isolation Kit (from Invitrogen). CD8 T cells were >99% pure and assessed by flow cytometry staining with CD8-Pacific Blue antibody (Biolegend). Total CD8 T cells were activated using anti-CD3/CD28 coated beads (Invitrogen) at a 1:2 ratio (beads:cells) and infected with anti-CD19 CAR, anti-Spike CAR or ACE2 CAR VSVG pseudotyped lentiviral constructs with multiplicity of infection (MOI) of 5-10. The cells were then expanded in complete RPMI 1640 medium supplemented with 10% Fetal Bovine Serum (FBS, Atlanta Biologicals), 1% penicillin/streptomycin (Corning Cellgro) and 20ng/ml of IL-2 and cultured at 37°C and 5% CO_2_ supplemented incubators. Respective viruses were added 24 hours after the activation. Cells were expanded for 10-12 days and cytotoxicity assays were performed following their expansion. To generate HEK-293T cells that transiently expressed wild type and mutated spike protein (ATCC; mycoplasma-free low passage stock), the cells were transfected with Spike protein expressing pLP plasmids using Lipofectamine 3000 (Invitrogen) according to the manufacturer’s protocol and stained for their spike protein expression 72 hours after the transfection as described in Staining and Flow cytometry Analysis. All engineered and wild-type HEK-293 and T2 cells were cultured in complete RPMI 1640 medium (RPMI 1640 supplemented with 10% FBS; Atlanta Biologicals, Lawrenceville, GA), 8% GlutaMAX (Life Technologies), 8% sodium pyruvate, 8% MEM vitamins, 8% MEM nonessential amino acid, and 1% penicillin/streptomycin (all from Corning Cellgro). To generate T2s and 293s with stable Spike overexpression, wild-type T2 and 293 cells were transduced with 3 MOI of Spike protein overexpressing VSVG lentivirus and proliferated. The infection levels were determined by GFP expression through Flow Cytometry analysis. For ACE2 overexpression in 293, wild-type ACE2 sequence was obtained from Ensembl Gene Browser (Transcript ID: ENST00000252519.8) and codon optimized with SnapGene by removing restriction enzyme recognition sites that are necessary for subsequent molecular cloning steps preserving the amino acid sequence, synthesized in GenScript and then cloned into a lentiviral vector. VSVG pseudotyped lentiviruses of respective constructs were generated as mentioned above and added to the cells with MOI of 3. Transduction levels were determined by ACE2 staining via Flow Cytometry 72 hours after the infection. ACE2 staining is described in Staining and flow cytometry analysis.

### Flow cytometry analysis

Cells were resuspended in staining buffer (PBS + 2% FBS) and incubated with fluorochrome-conjugated antibodies for 30 min at 4°C. CD8 T cells were identified with CD8-Pacific Blue antibody (Biolegend). Activation of CD8 CAR-T cells was determined with CD25 staining using CD25-APC antibody (Biolegend). CAR expressions of ACE2 CAR and anti-Spike CAR and ACE2 expression of ACE2-293 cells were determined with SARS-CoV-2 S1 protein, Mouse IgG2a Fc Tag (Acro Biosystems) incubation followed with APC Goat anti-mouse IgG2a Fc Antibody (Invitrogen) staining and RFP expression. CAR expression of anti-CD19 CAR was determined with Human CD19 (20-291) Protein, Fc Tag, low endotoxin (Super affinity) (Acro) followed by a secondary staining with APC conjugated anti-human IgG Fc Antibody (Biolegend) and RFP expression. For cytotoxicity assay analysis, stably Spike protein-expressing T2 and 293 cell lines were identified with GFP marker. For Spike protein flow cytometry analysis, the cells were stained with Biotinylated Human ACE2 / ACEH Protein, Fc, Avitag (Acro Biosystems), then stained with APC anti-human IgG Fc Antibody clone HP6017 (Biolegend). Samples were acquired on a BD *FACSymphony A5 analyzer* and data were analyzed using FlowJo (BD Biosciences).

### Cytotoxicity assay

Following the expansion of engineered CAR-T cells for 10-12 days, the cells were analyzed for their RFP and CAR expressions. Effector to target cell ratio was calculated based on the number of CAR expressing cells. CAR expressing cells were titrated from 2:1 to 1:16 effector to target cell ratio at 2-fold dilutions while the target cell number was constant. For ACE2-Bite cytotoxicity assays, resting total CD8 T cells were combined with wild-type Spike overexpressing 293 cells, empty vector transduced 293 cells, mutated Spike protein transfected 293 cells and wild-type 293 cells in a 4:1 Effector/Target cell ratio, and ACE2-Bite and control supernatant were added in 1:10 supernatant/cell medium ratio. Cytotoxicity assay conditions were analyzed with Flow Cytometry at 72 hours post-coculture and the cells were identified as described in Staining and flow cytometry analysis.

### ACE2-Bite detection assay

Supernatants from ACE2-Bite secreting and wild-type suspension 293 cells were collected at several timepoints with different cell densities ranging from 3 to 7 million/mL. ACE2-Bite molecules taken from 3 million/mL cell culture supernatant were concentrated 5-folds and 30-folds by using 15mL 30kDa MilliporeSigma™ Amicon™ Ultra Centrifugal Filter Units. To capture the ACE2-Bite or ACE2-Fc molecules, The DevScreen SAv Bead kit (Essen BioScience, MI) was used. Biotinylated 2019-nCoV (COVID-19) spike protein RBD, His, Avitag, Biotinylated SARS-CoV-2 Spike Trimer, His,Avitag™ (B.1.1.529/Omicron) (MALS verified) and Biotinylated SARS-CoV-2 S protein, His,Avitag™, Super stable trimer (MALS verified) were coated to SAv Beads according to manufacturer’s instructions. Confirmation of successful bead conjugation was determined by staining with anti-His Tag (Biolegend) and flow cytometry analysis. The conjugated beads were then used as capture beads in flow immunoassay where they were incubated with recombinant Human ACE2-Fc (Acro Biosystems) or ACE2-Bite supernatant samples for 1 h at room temperature. Supernatant samples were assayed at a 1:1 starting dilution and three additional tenfold serial dilutions. ACE2-Fc was tested at a 30 μg/mL starting concentration and in additional five threefold serial dilutions. Detection reagent was prepared using Human CD3 epsilon Protein, Mouse IgG2a Fc Tag (Acro) and Phycoerythrin-conjugated Goat anti-Mouse IgG2a Cross-Adsorbed Secondary Antibody (Fisher) for ACE2-Bite and APC anti-human IgG Fc Antibody clone HP6017 (Biolegend) for ACE2-Fc were added to the wells and incubated for another hour at room temperature. Plates were then washed twice with PBS and analyzed by flow cytometry using iQue Screener Plus (IntelliCyt, MI). Flow cytometry data were analyzed using FlowJo (BD biosciences). DevScreen SAv Beads were gated using FSC-H/SSC-H, and singlet beads gate was created using FSC-A/FSC-H. Gates for different DevScreen SAv Beads were determined based on their fluorescence signature on RL1-H/RL2-H plot (on iQue plus). PE fluorescence median, which is directly associated with each single plex beads was determined using BL2-H (on iQue plus). Geometric means of PE fluorescence in different titrations were used to generate the titration curve and the area under the curve was calculated using GraphPad Prism 9.0 software (GraphPad Software).

### Spike pseudotyped virus neutralization assay

Three-fold serially diluted recombinant human ACE2-Fc (Acro Biosystems) or two-fold serially diluted ACE2-Bite and control supernatants were incubated with GFP-encoding SARS-CoV-2 Spike pseudotyped viruses with 0.2 multiplicity of infection (MOI) for 1 hour at 37°C degrees. The mixtures were subsequently added to ACE2+ 293 cells, which were previously stained for their ACE2 expressions and confirmed ∼%100 positive before neutralization assays, for 72h hours after which cells were collected, washed with FACS buffer (1xPBS+2% FBS) and analyzed by flow cytometry using BD *FACSymphony A5 analyzer.* Cells that do not express GFP were used to define the boundaries between non-infected and infected cell populations. Percent infection was normalized for samples derived from cells infected with SARS-CoV-2 pseudotyped virus in the absence of ACE2-Fc or ACE2-Bite.

### Statistical Analyses and Reproducibility

All statistical analyses were performed, and graphs were prepared using GraphPad Prism V9 software. The numbers of repeats for each experiment were described in the associated figure legends.

## Results

### Development of SARS-CoV-2 specific synthetic CARs expressed in T cells

We sought to develop a system to test whether we can target cells that express SARS-CoV-2 Spike protein on their cell surface during the infection (16) (**Figure 1A**) employing effector human T cells engineered to express CAR molecules that can recognize the Spike protein on cell surface. For that, we transfected the cells with a plasmid containing a full-length wild-type Spike protein gene under CMV promoter (**Figure 1B**). 72 hours later, cells were stained with a recombinant ACE2-Fc protein followed by a fluorescent anti-Fc antibody to detect surface Spike protein expression and were compared to control cells which were transfected with Vesicular stomatitis Virus G (VSVG) plasmid. 293 cells transfected with full-length Spike protein plasmid displayed cell surface Spike expression indicating Spike protein indeed can be localized to the cell membrane despite its Endoplasmic Reticulum Retention Signal (ERRS) domain (**Figure 1C**). We then established a target 293 cell line that stably expressed Spike protein and Green Fluorescent Protein (GFP) as a reporter. Further, to enhance cell surface spike protein expression as shown in previous studies (17, 18), we deleted the ERRS domain. 72 hours after the transduction, engineered 293 cells were stained for their Spike expression, and flow cytometry analysis showed a co-expression of Spike and GFP in a high percentage of cells (**Figure 1D**).

**Figure 1:**
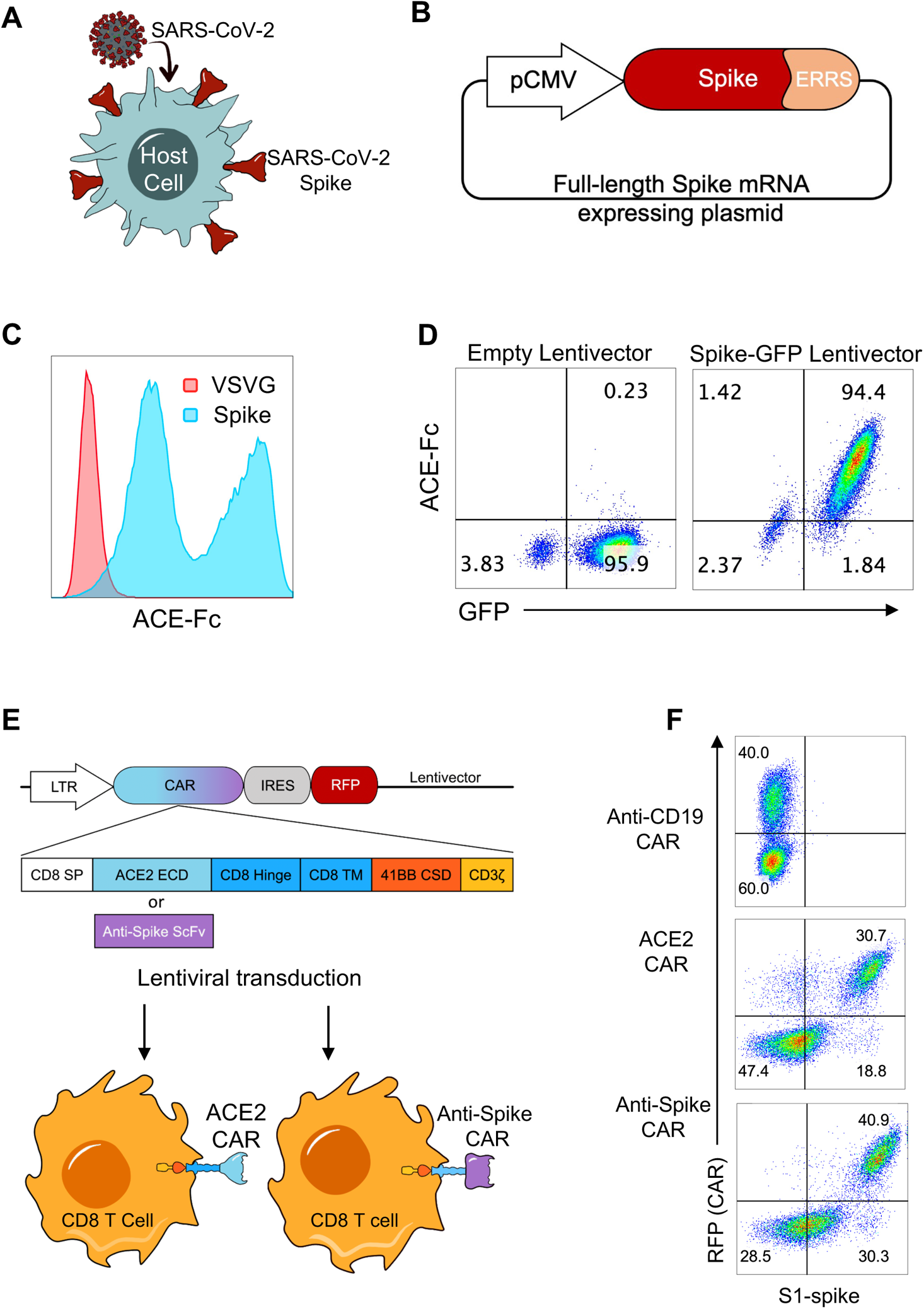
Engineering human primary CD8 T cells to express CAR molecules targeting SARS-CoV-2 Spike protein expressing cells. **(A)** Illustration of Spike protein localization on the surface of SARS-CoV-2 infected cells and **(B)** of full-length SARS-CoV-2 Spike protein expressing plasmid including the Endoplasmic Reticulum Retention Signal (ERRS) of Spike protein on C terminal. **(C)** 293 cells transfected with full-length Spike protein (blue histogram) or with VSV-G as a negative control (red histogram) expressing vectors. The cells were stained with ACE2-Fc and anti-Fc-APC secondary antibody, flow cytometry data overlays are shown. **(D)** 293 cells transduced with a lentivirus encoding a truncated Spike protein gene without the ERRS domain and Green Fluorescent Protein (GFP) as a reporter. Transduced cells were stained with ACE2-Fc and anti-Fc-APC secondary antibody, representative flow cytometry data plots are shown. **(E)** Illustration of ACE2 CAR and anti-SARS-CoV-2 Spike protein CAR constructs and their expression in CD8 T cells. A constitutive LTR promoter drives ACE2 or anti-Spike CAR and RFP genes separated by an Internal Ribosomal Entry Site (IRES). CAR constructs consist of CD8 alpha signal peptide, ACE2 or single chain variable fragment of an anti-Spike antibody, CD8 Hinge, CD8 transmembrane domain, 4-1BB (CD137) co-stimulatory domain and CD3ζ domain. Lentiviruses containing CARs were used to transduce primary CD8 T cells. **(F)** Expression of CAR constructs on CD8 T cells. Activated and transduced CD8 T cells were expanded for 10-12 days and stained with SARS-CoV-2 S1 protein fused to mouse Fc, and anti-mouse Fc secondary antibody. Flow cytometry plots showing ACE2 or anti-Spike surface expression versus RFP are shown. Anti-CD19 CAR expressing CD8 T cells were used as control. The experiments were replicated several times with similar results.

Next, we designed lentivector constructs containing ACE2 CAR or anti-Spike CAR cassettes followed by an Internal Ribosomal Entry Site (IRES) and Red Fluorescent Protein (RFP) and transduced human primary CD8 T cells (**Figure 1E**) as previously described (19). ACE2 CAR and anti-Spike CAR constructs comprised of CD8 alpha signal peptide, ACE2 extracellular domain (ECD) or anti-Spike ScFv, respectively, and intracellular 41BB co-stimulatory domain (CSD), and CD3ζ (zeta) signaling domains (**Figure 1E**). An Anti-CD19 CAR-RFP lentiviral construct was also designed to be used as a control. CD8 T cells were then activated and transduced with these lentiviruses encoding the CAR constructs and expanded in IL-2 for 10-12 days. Surface expression of ACE2 and anti-Spike was demonstrated on CAR engineered CD8 T cells, which also correlated with RFP reporter (**Figure 1F**).

### Cytotoxicity assays with ACE2 CAR and anti-Spike CAR expressing T cells

We then co-cultured Spike+ target cell line and effector T cells expressing CARs and measured the cytotoxic activity of the T cells (**Figure 2A**). Briefly, after the ∼2-week proliferation of CAR-T cells, the cells were co-cultured for 72 hours with Spike-expressing target cells at different effector to target ratios. The CD8 T cells were then stained with anti-CD25 to determine their activation. Target cells were identified via GFP, which was co-expressed with Spike protein. Both ACE2 CAR and anti-Spike CAR-T cells became highly activated and killed the Spike+ 293 cells whereas control anti-CD19 CAR-T cells were neither activated nor showed any cytotoxicity (**Figure 2B, 2C**).

**Figure 2:**
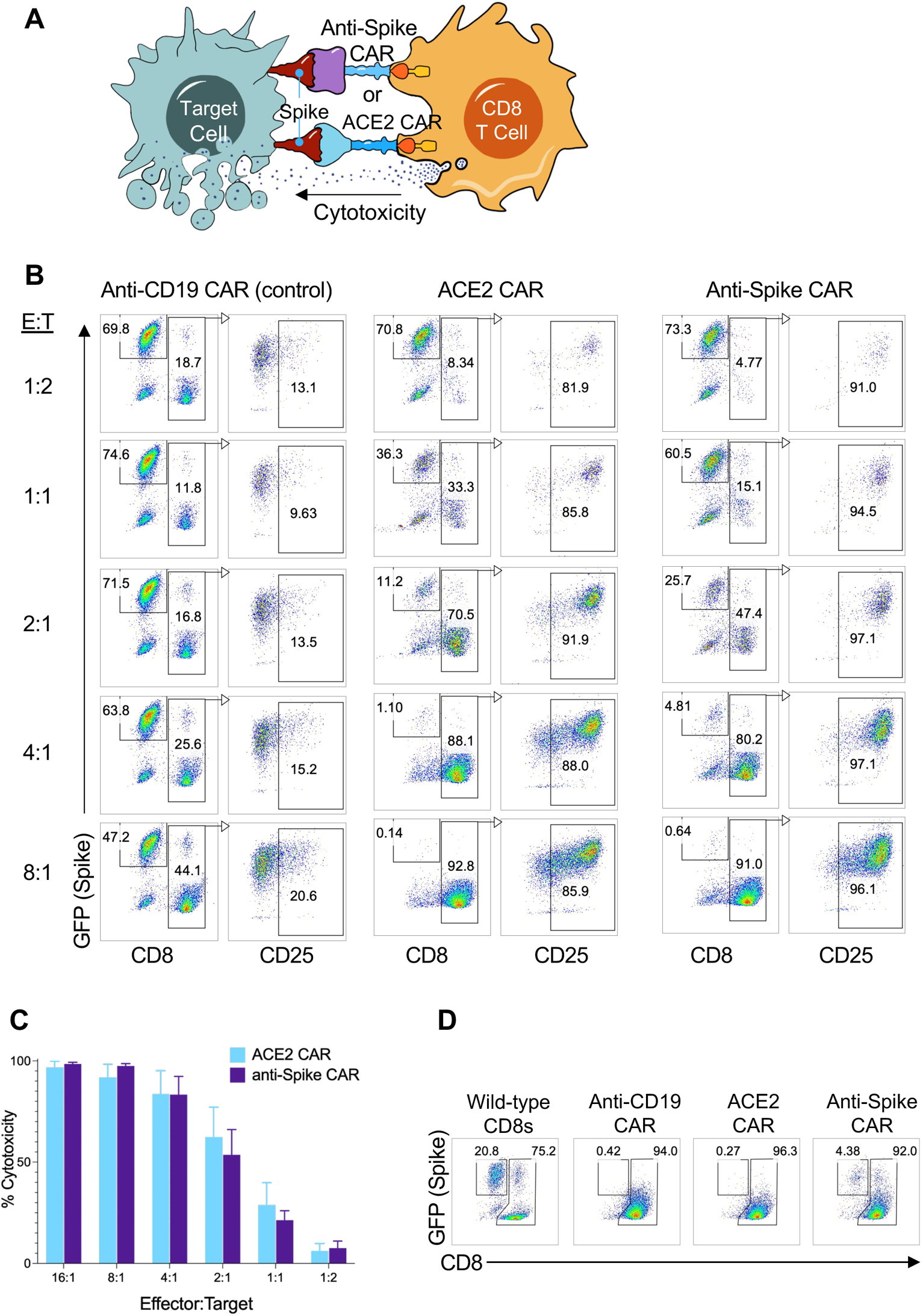
Cytotoxic activity of human primary CD8 T cells engineered to express ACE2 CAR or anti-Spike CAR. **(A)** Illustration of cytotoxicity assay against Spike-expressing target cells using ACE2 CAR or anti-Spike CAR expressing CD8 T cells as effector cells. **(B)** CAR engineered T cells cytotoxicity assays with Spike expressing 293 target cells at different Effector:Target ratios. CD8 T cells transduced with anti-CD19 CAR lentiviruses were used as control effector cells. Effector CD8 T cells were identified with CD8 staining while target cells were gated based on GFP (Spike) expression. Activation of effector cells and CAR expression were determined with CD25 expression after gating on CD8 T cells 2 days after co-culture. **(C)** Percent cytotoxicity of ACE2 CAR (blue) and anti-Spike CAR (purple) T cells normalized to anti-CD19 CAR-T cells at different Effector:Target ratios and using Spike-expressing 293 cells as the target. **(D)** CAR engineered T cells cytotoxicity assays with Spike-expressing target B cell line (T2 cells) at 8:1 E:T ratio. Wild-type CD8 T cells were used as negative control and anti-CD19 CAR expressing CD8 T cells were used as positive control. Panels show representative experiments replicated with similar results.

We next tested whether ACE2 CAR and anti-Spike CAR-T cells can kill Spike-expressing human B cell line, which can also be used as positive control using anti-CD19 CAR-T cells. ACE2 CAR and anti-Spike CAR-T cells killed Spike-expressing B cells as efficiently as 293 cells, indicating that different cell types infected with SARS-CoV-2 can be targeted using these novel CAR-T cells (**Figure 2D**). In addition, ACE2 CAR and anti-Spike CAR-T cells did not show cytotoxicity to GFP-expressing, Spike-negative control targets and were also not activated, showing a selective Spike protein-mediated activation and killing (**Supplementary Figure 1**).

### Development of bispecific antibodies to mobilize and activate T cells against SARS-CoV-2 Spike protein-expressing target cells

Currently, the CAR-T cell immunotherapy procedure requires a meticulous process of collecting cells from patients, engineering them in a good manufacturing process environment, re-infusion, and extensive clinical follow-up of the patients (20). As such this may not be very practical for treatment of COVID-19 patients. To overcome these hurdles of CAR-T approach, we engineered bispecific antibodies (Bites) as T cell activators, consisting of an anti-CD3 scFv fused with the extracellular domain of ACE2 to redirect CD3 T cells to SARS-CoV-2 infected cells (**Figure 3A**). The ACE2-Bite cassette consisted of ACE2 signal peptide, ACE2 extracellular domain, a linker peptide, an anti-CD3 antibody single-chain variable fragment, a His-Tag, and a Hemagglutinin (HA) Tag (**Figure 3B**). ACE2-Bite was produced by suspension 293 cells as described in methods. The supernatant from these cells were then filtered to eliminate small molecules, which also resulted in ∼30-fold concentration of ACE2-Bite proteins. The supernatant of wild-type suspension 293 cells was also collected and filtered/concentrated to be used as a control.

**Figure 3:**
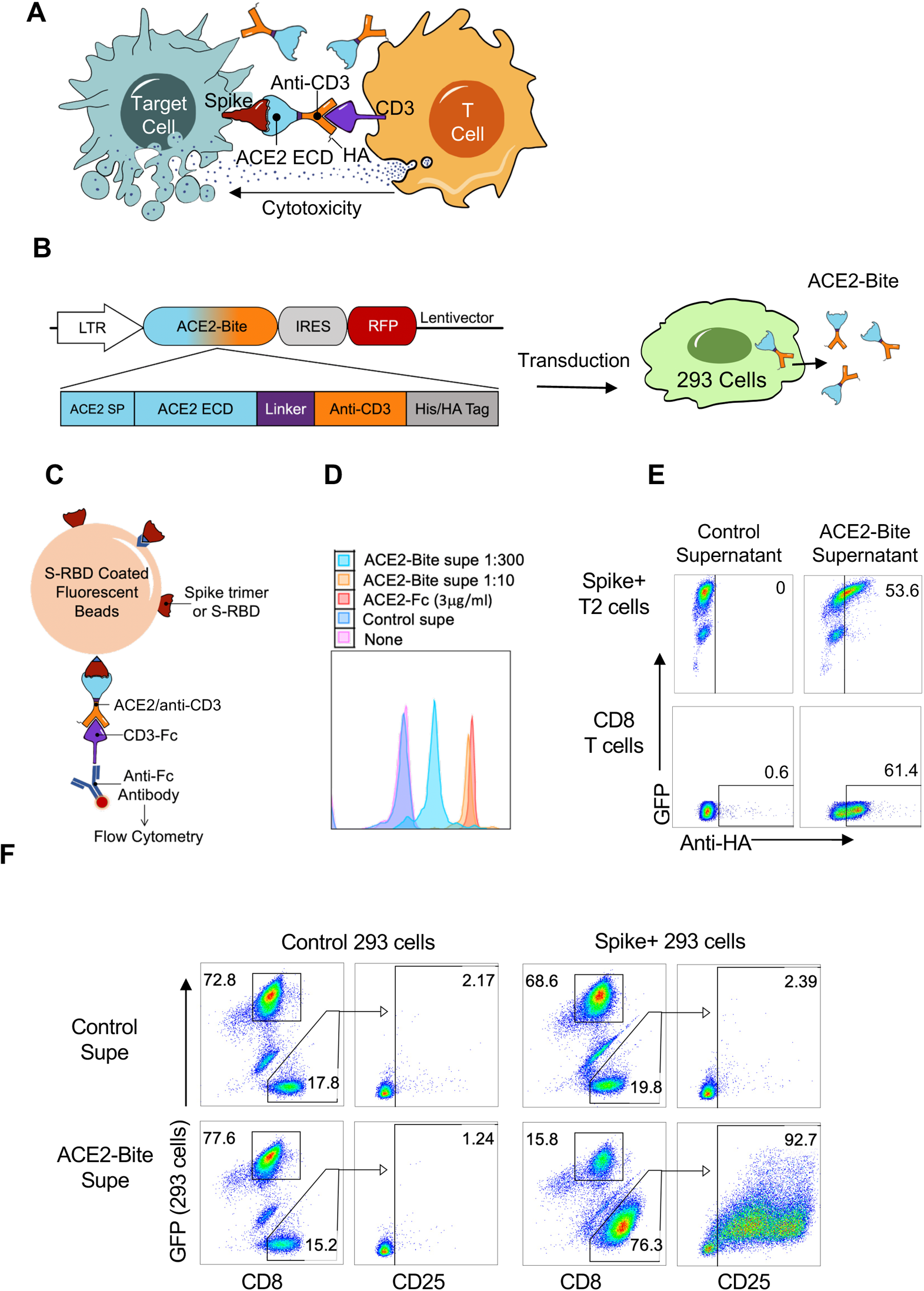
Functional ACE2/anti-CD3 bi-specific T cell engagers against SARS-CoV-2. **(A)**. Illustration describing mechanism of action of ACE2-Bite. The extracellular domain (ECD) of ACE2 (blue) in ACE2-Bite binds to Spike protein (red) expressed on the surface of SARS-CoV-2 infected cells and the anti-CD3 fragment (orange) binds to CD3 molecule (purple) on T cells linking both cell types and inducing the activation of T cells which subsequently results in apoptosis of infected target cells. ACE2-Bite recombinant protein also contains a hemagglutinin (HA) tag at the C terminal. **(B)** Representation of ACE2-Bite construct and the protein production in 293 cells. A constitutive LTR promoter drives the expression of ACE2-Bite and RFP genes separated by an Internal Ribosomal Entry Site (IRES). ACE2-Bite cassette consists of ACE2 signal peptide (SP), ACE2 extracellular domain, a linker peptide, an anti-CD3 antibody single-chain variable fragment, a His-Tag, and a Hemagglutinin (HA) Tag. Lentiviruses expressing ACE2-Bite were used to transduce suspension 293 cells that produce and secrete ACE2-Bite protein in their culture supernatant. **(C)** Illustration of the bead-based ACE2-Bite capture assay. Fluorescent beads coated with Spike protein trimer or Spike-Receptor binding domain (S-RBD) were used to capture ACE2-Bite molecules which were detected via a recombinant CD3-Fc fusion protein and an anti-Fc antibody then subsequently analyzed by flow cytometry. ACE2-Fc molecules were also detected with Spike trimer or S-RBD coated beads and anti-Fc antibody**. (D)** Detection of different concentrations of ACE2-Bite (1:10 and 1:300 dilutions were shown in orange and turquoise, respectively) and ACE2-Fc (3 μg/mL) (red) using bead-based ACE2-Bite capture assay. Wild-type 293 cell supernatant (Control supe, Blue) and staining buffer (None, Pink) were used as negative controls. **(E)** Binding of ACE2-Bite to Spike-GFP expressing T2 cell line and primary human T cells. HA staining of Spike-GFP expressing T2 cells (top panel) and CD8 T cells (bottom panel) when combined with ACE2-Bite (right plot) or control (non-transduced 293) (left plot) supernatants. **(F)** CD25 and GFP expressions show activation and cytotoxicity of resting CD8 T cells against Spike/GFP-expressing or control (transduced with GFP-expressing empty vector) 293 cells in the presence of ACE2-Bite or control supernatant. The experiments were replicated with similar results.

To test the correct folding of the recombinant ACE2-Bite protein, we developed a fluorescent bead-based ACE2-Bite detection assay in which the fluorescent beads were coated with either SARS-CoV-2 Spike protein trimer or Spike-Receptor Binding Domain (S-RBD) and the ACE2-Bite molecules captured by these beads were detected via a recombinant CD3-Fc molecule which was then stained with an anti-Fc antibody (**Figure 3C**). A recombinant ACE2-Fc molecule was used as a positive control since ACE2 part could bind to Spike trimer or S-RBD on the surface of beads and anti-Fc antibody could recognize the Fc part of ACE2-Fc. ACE2-Bite detection assay with S-RBD coated beads showed that secreted and concentrated ACE2-Bite levels (1:10) were comparable to control ACE2-Fc concentration (3 μg/mL) (**Figure 3D**). We also confirmed that ACE2-Bite concentration protocol functioned as intended and increased the ACE2-Bite concentration by an order of magnitude **(Supplementary Figure 2)**.

We then tested the ACE2-Bite binding on human primary CD8 T and Spike-expressing target cells. For this, ACE2-Bite and wild-type supernatants were added to primary human CD8 T cells and a B cell line (T2 cells) which was engineered to express Spike/GFP. The cells combined with ACE2-Bite or control supernatants were then stained for HA Tag on their surface. Spike/GFP co-expressing T2 cells and CD3 expressing primary human CD8 T cells combined with ACE2-Bites were stained positive for HA Tag, suggesting Spike-specific binding of ACE2 fragment and CD3-specific binding of Anti-CD3 fragment. (**Figure 3E**).

We then performed a cytotoxicity assay to test the ability of ACE2-Bites to trigger human T cell activation and effector function. Human primary CD8 T cells were co-cultured with Spike-expressing or control 293 cells in the presence of ACE2-Bite or control supernatants. 2 days later cells were collected and stained for their CD8 and CD25 expression. GFP expressed by control and Spike lentivectors was used to identify the target cells. Indeed, resting human T cells became activated and were cytotoxic only in the presence of ACE2-Bite supernatant and Spike-expressing targets, suggesting Spike-specific T-cell activation functionality of ACE2-Bites (**Figure 3F**).

### Determining function of ACE2-Bite on Spike protein variants

A major advantage of targeting the SARS-CoV-2 Spike protein through its receptor ACE2 using ACE2-Bite is that this approach is less affected by antibody escape mutations, as mutated Spike proteins would still need to interact with ACE2. In fact, it is conceivable that variants with increased affinity to ACE2 would bind better to ACE2-Bite, possibly improving its efficacy.

To test this with an ACE2-Bite/Spike affinity assay, we first employed our bead-based antibody detection strategy described in Figure 3. In this experiment, we labelled the beads with wild-type and Omicron Spike protein trimers to determine the affinity of ACE2-Bites to natural trimeric Spike protein structure of these strains. Labelled beads were treated with ACE2-Bite and control supernatants in three-fold serial dilutions from 1 to 30 to generate titration curves. ACE2-Bite treated beads were then stained with recombinant CD3-Fc protein and anti-Fc antibody and analyzed via flow cytometry. Bead-based assay using these Spike protein trimers revealed a significant increase in affinity of ACE2-Bite molecule to Omicron Spike trimer compared to wild-type (p<0.0001) (**Figures 4A, B**).

**Figure 4:**
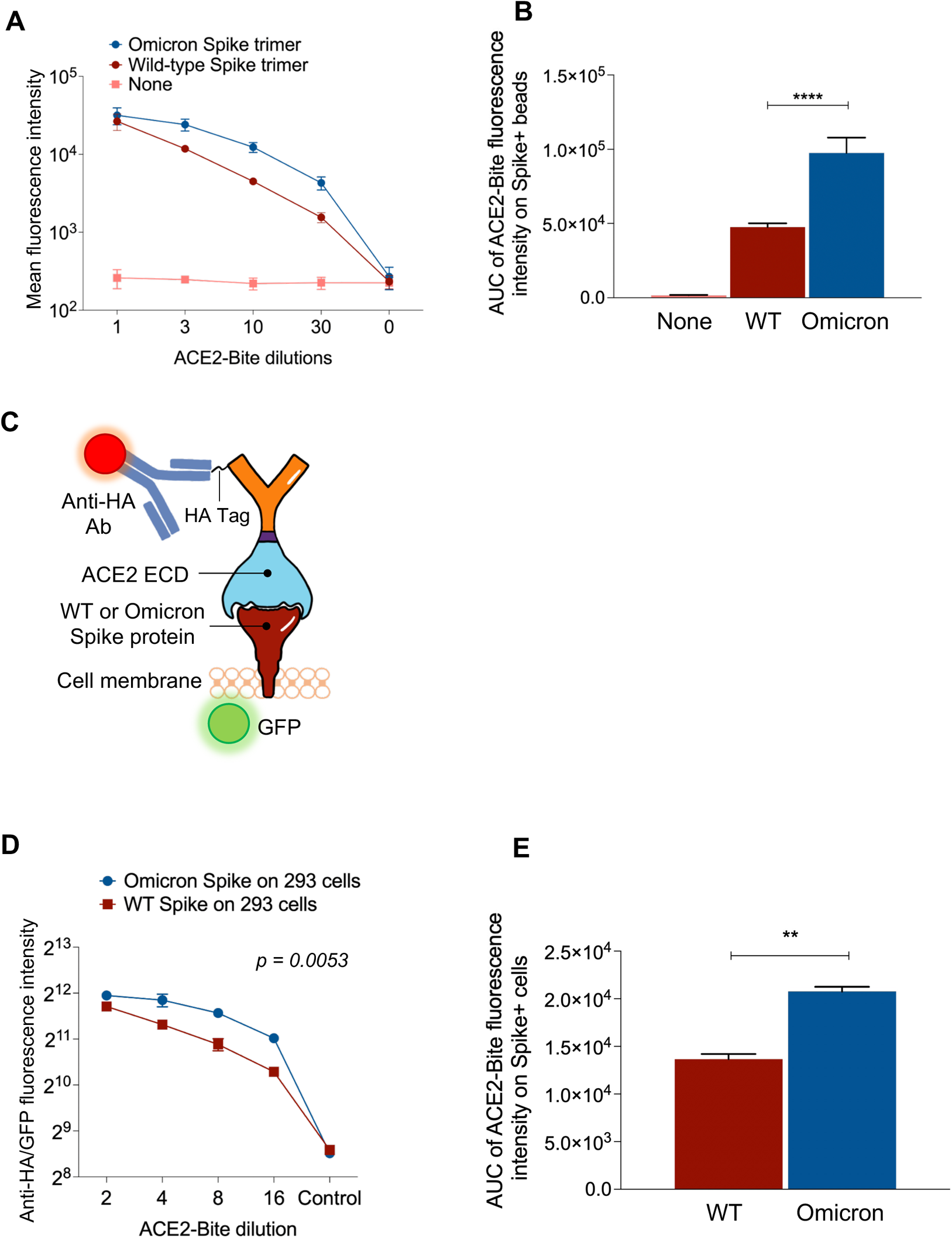
Binding of ACE2-Bite to variant Spike proteins. **(A)** Bead-based ACE2-Bite/Spike binding assay. Beads were coated with full length SARS-CoV-2 wild-type and omicron variant Spike protein trimers and treated with ACE2-Bite and control media in three-fold serial dilutions from 1 to 30. Treated beads were then stained with a CD3-Fc recombinant protein and an anti-Fc antibody. Geometric mean values of anti-Fc antibody fluorescence in flow cytometry were used to quantify the fluorescent intensity of samples. **(B)** Area under the curve (AUC) values of bead-based ACE2-Bite/Spike binding assay data from Figure 4A. Experiments were replicated three times. **(C)** Illustration of ACE2-Bite binding to spike protein (wild-type or Omicron) expressed on the cell surface and its detection by immunostaining with an anti-HA antibody. GFP was co-expressed with Spike proteins as a reporter. **(D)** ACE2-Bite binding assay on Spike-expressing 293 cells. Geometric mean intensity of anti-HA antibody staining used to quantify the affinity of ACE2-Bite molecules on Spike-expressing cells. For each condition, gates with similar intensities of GFP used as a Spike protein marker when assessing the fluorescence intensity of ACE2-Bite-stained cells to determine the quantitative value of ACE2-Bite fluorescence per Spike protein. Fluorescence intensity was determined via geometric mean values. Unpaired t test was used to determine the statistical significance and p values were corrected for multiple comparisons using Holm-Sidak method. **(E)** Area under the curve (AUC) values of bead-based ACE2-Bite/Spike binding assay data from Figure 4D. The experiments were replicated twice with similar results.

In addition to performing affinity assay on Spike-labeled beads, we also transduced 293 cells with GFP encoding lentivectors to express wild-type and Omicron variant Spikes and determined the ACE2-Bite affinity to these proteins on live target cells (**Figure 4C**). 3 days after the transduction, the cells were collected and co-stained with ACE2-Bite and anti-HA antibody and analyzed via flow cytometry. The ACE2-Bite/Spike protein binding assay on living cells revealed a significantly higher affinity of ACE2-Bite to Omicron variant Spike protein compared to Wild-type (p=0.0053), confirming the result of the bead-based affinity assay (**Figures 4D, E**). Taken together, these results implicate the potential pan-SARS-CoV-2 effectivity of ACE2-Bite approach, even further, possibly higher efficiency of the neutralization and T cell cytotoxicity, as the virus evolves into variants with higher affinity towards ACE2 protein.

### ACE2-Bite neutralizes Spike-pseudotyped lentiviruses

In addition to bridging infected cells to activate T cells, we reasoned that ACE2-Bite may also neutralize SARS-CoV-2 by binding to Spike proteins on the virus. To test this, we generated a set of lentiviruses pseudotyped with SARS-CoV, SARS-CoV-2 wild-type, Delta, and Omicron Spike proteins and investigated ACE2-Bite neutralization on these viruses. Neutralization assay was performed by pre-culturing pseudotyped viruses with different dilutions of ACE2-Bite supernatant and then adding to ACE2 expressing 293 cells as previously described (21) (**Figure 5A**). We also incubated a recombinant ACE2-Fc molecule at different concentrations with Spike pseudotyped lentivirus as a positive control. The infection levels were determined 3 days post-infection based on the GFP expression of ACE2-expressing 293 cells. As shown in the representative experiment, ACE2-Fc and ACE2-Bite molecules neutralized the Spike pseudotyped lentivirus (**Supplementary Figure 3**). Importantly, ACE2-Bite molecule was able to neutralize all versions of Spike encoding lentiviruses with increased efficiency against Delta and Omicron variants compared to wild-type SARS-CoV-2 pseudovirus (p=0.016, 0.0008 and 0.016 for Delta 1:28, 1:14 and 1:7 dilutions, respectively; p=0.0023 and 0.014 for Omicron 1:14 and 1:7 dilutions, respectively) (**Figure 5B**). These neutralization assays demonstrated that novel ACE2-Bite recombinant protein could also function as a decoy receptor for the virus and could perform better against new emerging variants. Taken together, these findings demonstrates the potential of ACE2-Bite as a fool-proof therapeutic approach.

**Figure 5:**
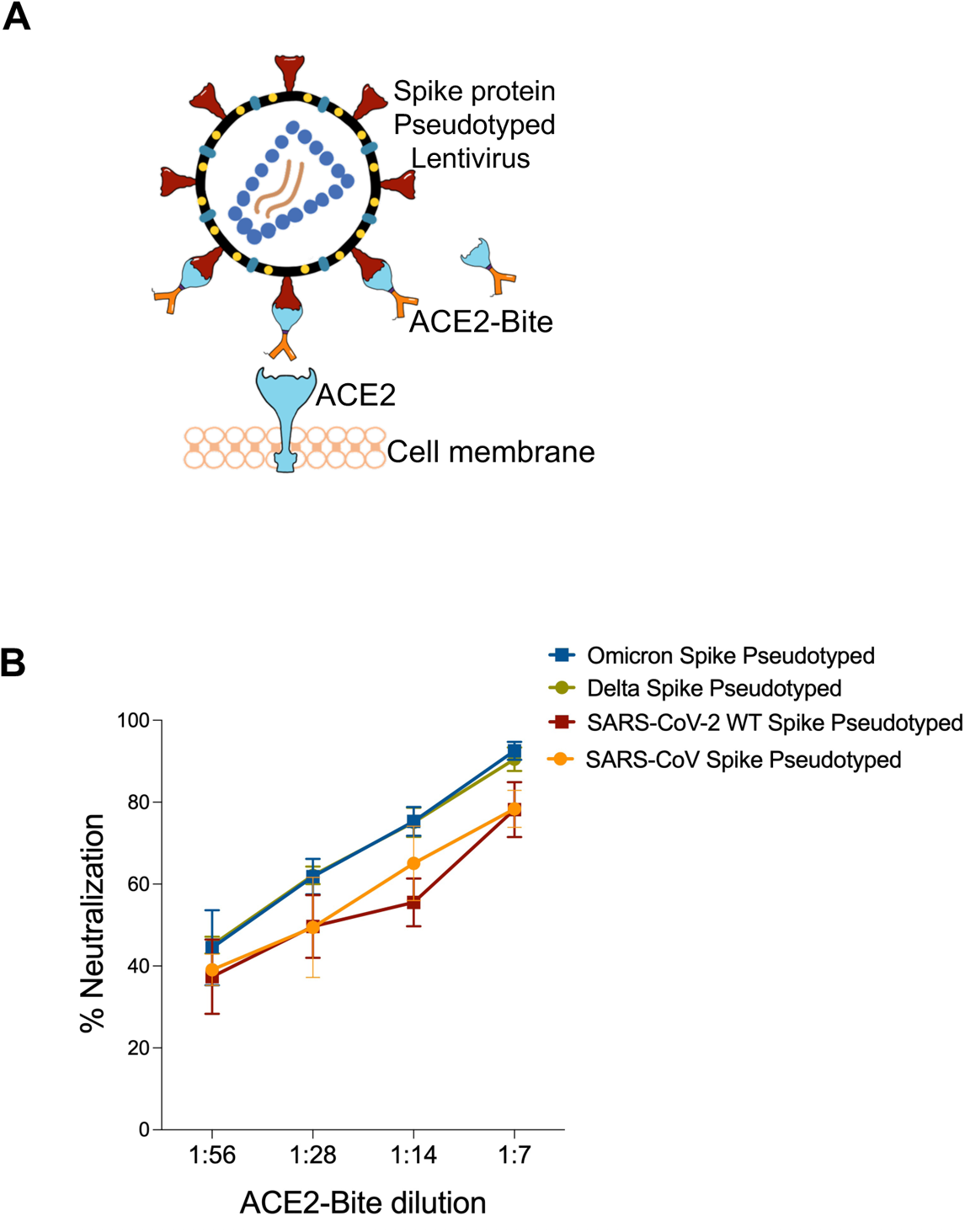
Binding of ACE2-Bite to SARS-CoV-2 Spike protein variants on pseudotyped lentiviruses for virus neutralization. **(A)** Schematic illustration of virus neutralization assay. ACE2-Bite and Spike (SARS-CoV, SARS-CoV-2, Delta and Omicron) pseudotyped lentiviruses are pre-incubated then added to ACE2-overexpressing 293 cells. **(B)** Line graph represents virus neutralization data of the lentiviruses pseudotyped with different Spike proteins that pre-incubated with ACE2-Bite at different ACE2-Bite:virus ratios then added to ACE2-overexpressing 293s. The experiments were replicated twice with similar results.

## Discussion

Despite advances in vaccine development, COVID-19 is still a major cause of morbidity and mortality in the USA and throughout the world. Rapid evolution of the virus is also a major concern, suggesting the need to develop novel effective treatment strategies, as SARS-CoV-2 specific targeted therapeutic approaches such as monoclonal antibody therapies can lose their effectiveness as new escape variants emerge. To mitigate or override these potential problems, here we utilized synthetic biology approaches, to develop synthetic molecules that can bridge T cells with SARS-CoV-2 infected cells, through recognition of cell surface expression of virus Spike protein and eliminate them through cytotoxic activity.

We first generated CD8 T cells expressing chimeric antigen receptors (CARs) specific to Spike protein with an anti-Spike antibody or ACE2 surface domain on the extracellular region and tested their effectiveness against different cell types expressing Spike proteins. In this assay, engineered CAR-T cells (anti-Spike CARs and ACE2 CARs) became activated and killed the Spike-expressing target cells selectively. Although cancer cells have predominantly been the focus of adaptive cellular immunotherapies, studies have suggested that autoimmune and infectious diseases could also be targeted via such approaches (22, 23). Several studies have employed or investigated CAR-T approach against infectious diseases, such as Human immunodeficiency virus (HIV) (24–26), Hepatitis B virus (HBV) (27), Hepatitis C virus (HCV) (28), Cytomegalovirus (CMV) (29), Epstein-Barr virus (EBV) (30) and Aspergillus fumigatus (31). While it is not practical to apply this approach during COVID-19, it is at least conceptually possible to develop CAR-T cells engineered to target infected cells as prophylaxis, in immune-compromised or elderly individuals who are in the high-risk category for mortality, as the lifespan of these cells could be many months to years. It may also be possible to generate off-the-shelf Spike-specific CAR-T cells by removing genes in T cells that can cause allogeneic reaction or graft versus host disease (32). This may also be an important proof-of-principle for other viral infections, current or future for which we do not yet have vaccines available (33, 34).

While Spike specific CAR-T cell demonstrated a proof-of-concept for directing T cells towards Spike expressing cells, this approach would not be practical in a clinical setting. As such, we developed bispecific T cell engagers (Bites), in protein forms, with the potential to activate cytotoxic cells upon bridging with Spike protein expressed on infected cell surface. We demonstrated that ACE2-Bite bispecific antibody-receptor complex, triggered effective CD8 T cell activation, which resulted in selective killing of Spike+ target cells. Indeed, a B cell surface protein (CD19) specific Bite called Blinatumomab (CD19-CD3 Bite) have already been approved to be used in B cell lymphoma patients in 2018 (35). Other studies also showed that Bites could be employed against Her2 (36), BCMA (37), EpCAM (38), EGFR (39), CD20 (40), and PDL1 (41) expressing cancers; and diseased cells infected by CMV (42) and HIV (43). Compared to current treatments (such as neutralizing antibodies or anti-virals) ACE2-Bite approach may potentially be effective both at early stages (as neutralizer of the virus entry) and later stages of the infection when antibody immune defenses are breached, and T cells become more important in restricting the spread of the virus in vivo. Another major advantage of the ACE2-Bite approach is using ACE2, the key host receptor of SARS-CoV-2, as the Spike protein recognizing part of the bi-specific antibody. As shown in **Figure 4**, this allowed us to target mutated Spike proteins from variants of concern with even better efficiency in contrast to conventional antibody approaches which lose their efficiency due to immune escape mutations (13).

In addition to recognizing mutated Spike proteins, we found that ACE2 part of the ACE2-Bite functioned as a decoy receptor and neutralized the virus, preventing it from infecting the cells. The neutralization feature of the ACE2-Bite molecule is promising for its use as preventive treatment and it would conceivably have synergistic effect with the cytotoxic effect by engaging T cells towards infected cells. Indeed, in line with our findings, a clinical study by Zoufaly *et al.* found that infusion of soluble recombinant human ACE2 molecule in a 45-year-old COVID-19 patient resulted in dramatic decrease in viral copies in the patient plasma (44). Other studies also demonstrated the neutralization capacity of soluble ACE2 molecule (45–47). Considering the immunity of ACE2 to the Spike mutations, both as a CD3 T cell redirecting molecule and a decoy receptor, the efficacy of the ACE2-Bite treatment is unlikely to be diminished by variants arising during COVID-19 pandemic or in possible future SARS pandemics.

If these approaches are developed as therapeutics, it will also be important to consider side effects of CAR-T cells or ACE2-Bite treatment, cytokine release syndrome and T cell exhaustion, similar to cancer focused CAR-T therapies (48). Although, we think this will be less likely given the number of infected cells would be expected to be much smaller compared to tumor burden; overall, much fewer T cells would be stimulated during the infection. Another potential downside could be that the ACE2 in ACE2-Bite may interact with its physiologic ligands and interfere with the renin-angiotensin system, although, so far recombinant human ACE2 molecule has been tested in 89 patients with tolerable clinical outcomes (49, 50). Regardless, if this becomes a problem it can be prevented by mutating the carboxypeptidase activity of ACE2. Furthermore, downregulation of ACE2 levels due to SARS-CoV-2 binding during COVID-19 is associated with hyperactivation of renin-angiotensin system and injuries to the lung, kidneys, and heart (51). Thus, ACE2-Bite may even have beneficial properties as a replacement to wild-type ACE2 activity during COVID-19.

In conclusion, engineered CD8 T cells expressing Spike protein-specific chimeric antigen receptors and ACE2/anti-CD3 bispecific T cell engagers developed in this study could be used to target SARS-CoV-2 infected host cells and the virus itself, and may be alternative future therapeutic strategies for COVID-19.

## Author Contributions

M.D., L.K. and D.U. conceived and designed the experiments. M.D., L.K., L.P., F.K., and M.Y. carried out the experiments. M.D., L.K., and D.U. analyzed the data, drew illustrations, and prepared the figures. M.D. and D.U. wrote the manuscript.

## Declaration of Interests

M.D. and D.U. are inventors in a provisional patent application.

## Acknowledgments

The research in this study was supported by National Institute of Health (NIH) grant U19 AI142733-01 (DU) and Achelis and Bodman Foundation (DU). We thank Courtney Gunter and Sara Cassidy for critical reading and Mustafa Semih Elitok for advice on designing anti-Spike antibody ScFv protein.

## Data and Material Availability

The source data for the Figures along with the Supplementary Figures are available upon request. All unique/stable reagents generated in this study are available from the lead contact with a completed materials transfer agreement

## Supplemental Figures

**Supplementary Figure 1:**
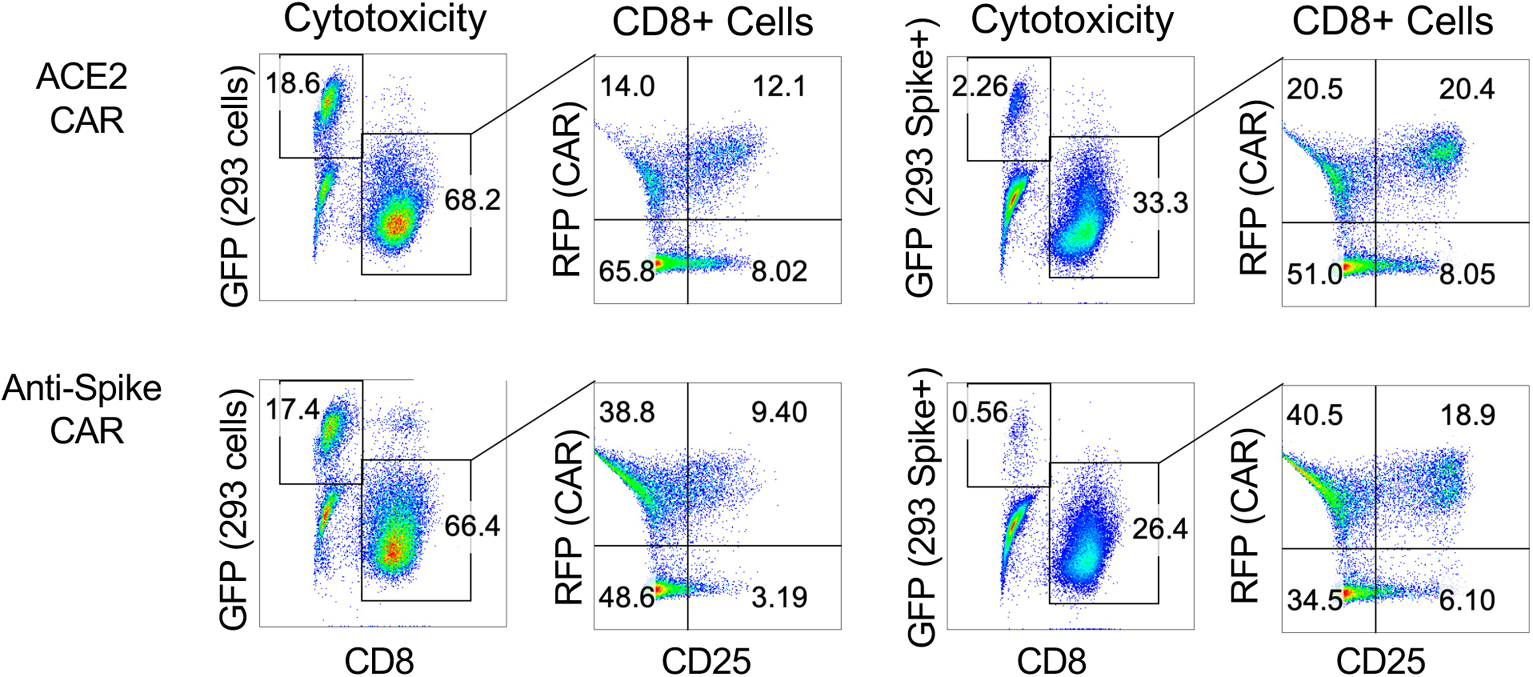
Selective cytotoxicity of ACE2 CAR and anti-Spike CAR-T cells for spike expressing target cells. CAR engineered T cell cytotoxicity assays with Spike expressing and control 293 target cells. Control 293s were engineered with a GFP-expressing empty vector. Spike-expressing and control 293s were identified by GFP expression. Effector cells were identified by CD8 staining. T cell activation was determined via CD25 staining. CAR expressing T cells co-expressed RFP with CAR constructs.

**Supplementary Figure 2:**
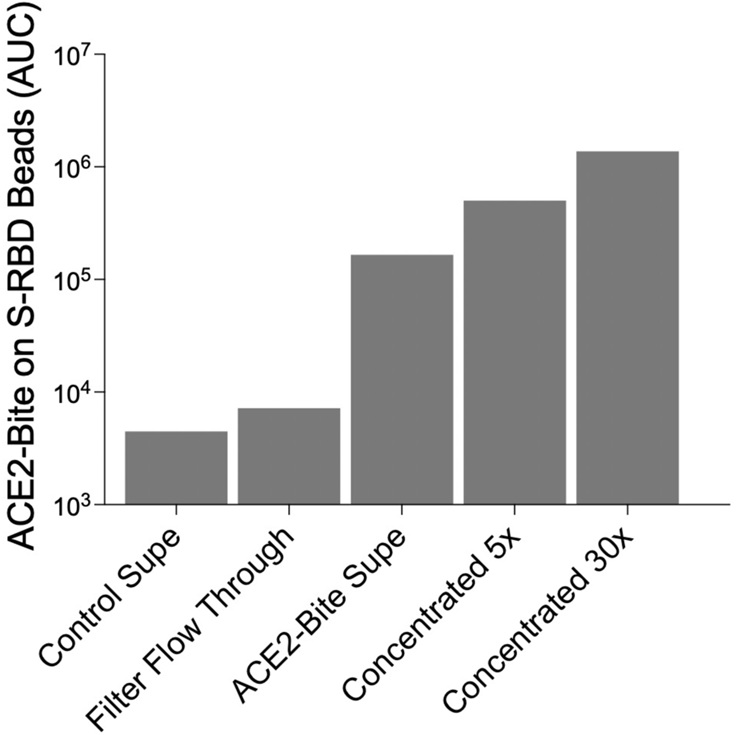
ACE2-Bite concentrations determined by a bead-based capture assay. Area under the curve (AUC) values of ACE2-Bite molecules in supernatants from different conditions. ACE2-Bite supernatant was concentrated 5-folds and 30-folds. Flow through supernatant from the concentration process (Filter flow through) and wild-type control supernatant were used as controls. Fluorescent beads coated with Spike-Receptor binding domain (S-RBD) were used to capture ACE2-Bite molecules in supernatants titrated from 1:1 to 1:1000 by 10-fold serial dilutions were detected via a recombinant CD3-Fc fusion protein and an anti-Fc antibody. Geometric mean of anti-Fc antibody fluorescence was used to generate the curves which were used to calculate the area under the curve values.

**Supplementary Figure 3:**
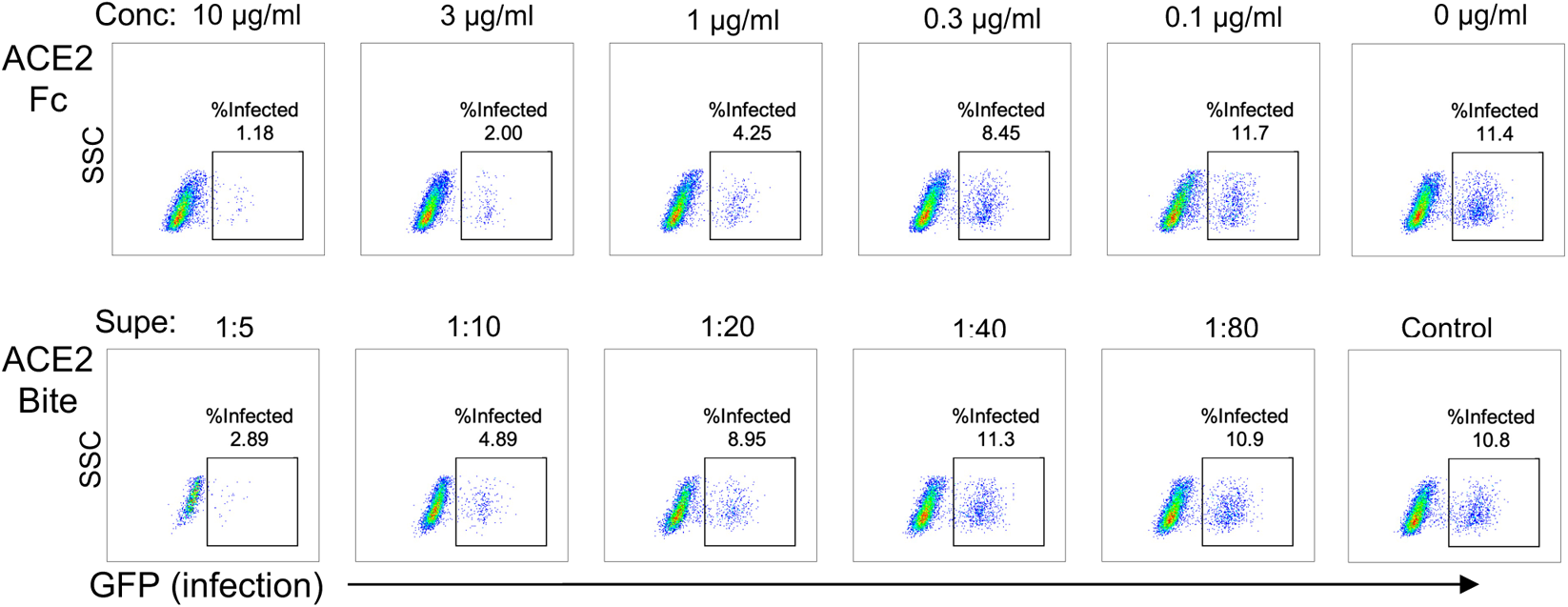
Neutralization efficiency of ACE2-Bite compared to recombinant ACE2-Fc protein. FACS plots show neutralization data of delta variant spike protein pseudotyped virus infection when pre-incubated with different concentrations of ACE2-Fc (top panel) or different dilutions of ACE2-Bite supernatant (bottom panel). The infection levels were determined 3 days later via flow cytometry based on GFP expression.

## Notes

### Competing Interest Statement

The authors have declared no competing interest.

### Summary of Updates

We revised the Title and abstract, and made some editing throughout the manuscript to clarify our points. A new figure 5 was added and figure 4 was edited, based on new data showing our approach inhibits Omicron variant better compared to wild type virus strains.

